# Magnetic field-based actuator-sensor assembly for dynamic mechanical loading and functional analysis of engineered heart tissues in culture

**DOI:** 10.64898/2026.11.12.766088

**Authors:** Nikolay Naumenko, Tomi Tuomainen, Robert S. Leigh, Pasi Tavi

## Abstract

Cultured *ex vivo* and stem cell-derived cardiac tissues have emerged as promising tools for discovering mechanisms of human disease, for pharmacological testing and for elucidating mechanisms of genetic cardiac diseases. The current culture systems for these tissues do not fully recapitulate the mechanobiological environment of cardiac tissue or allow detailed characterization of muscle function in culture, which limits the interpretation of these models. To address this methodological gap, we present an integrated electromagnetic actuator chamber (EMC) that modulates preload and afterload independently in engineered heart tissue (EHT) in culture. Because the load can be adjusted online, the system enables longitudinal monitoring of passive mechanical properties, including viscosity and elasticity. Combining these tissue-specific properties with measurement of muscle displacement by high-sensitivity magnetic field detectors and pre-calibrated electromagnetic force, the system provides simultaneous data on displacement and absolute contractile force. With independent modulation of preload and afterload, it enables switching from auxotonic to isometric contraction and determination of length-dependent activation, and these can be tracked over time during culture. To our knowledge, no earlier system combines independent, freely adjustable control of both preload and afterload with continuous, calibrated functional readout in culture, making it possible to study how mechanical load shapes cardiac tissue function, maturation and disease development.

## Introduction

Engineered heart tissues (EHTs) and other stem-cell-derived human cardiac models have emerged as powerful platforms for investigating human cardiac physiology and disease, and they appear increasingly well suited to drug discovery and safety testing (Anand et al., 2025; Campostrini et al., 2021). These models provide experimental access to human cardiac tissue with known genetic identity under controlled conditions and have thus enabled the investigation of mechanisms associated with human genetic cardiac diseases (Carvajal-Vergara et al., 2010; Itzhaki et al., 2011; Yazawa et al., 2011). Their broader translational potential, however, remains limited by incomplete tissue maturation and by the difficulty of reproducing and quantitatively interrogating the physical environment in which cardiac muscle normally functions. Consequently, increasing efforts have focused not only on improving the structural and functional maturity of engineered myocardium (Feyen et al., 2020; Hirt et al., 2014; Leonard et al., 2018; Min et al., 2024; Parikh et al., 2017), but also on developing approaches that more comprehensively recapitulate the working environment of cardiac tissue and characterize its mechanical function (Bliley et al., 2021; Burattini et al., 2024; Kurashina et al., 2023; Mannhardt et al., 2017; Mannhardt et al., 2019; Tsan et al., 2021).

Cardiac muscle rarely, if ever, operates under static mechanical conditions *in vivo*. In the heart ventricle, the force that stretches the muscle before contraction, set by diastolic filling, forms the preload, which in turn determines the level of force through length-dependent activation of the myofilaments. At the start of left ventricular contraction, the muscle contracts at fixed length, i.e. isometrically. After the aortic valve opens, the muscle shortens at variable force (auxotonic contraction) against the pressure in the arteries, which forms the afterload. Accordingly, cardiac function inherently consists of the interaction between active contractility, passive tissue mechanics, and dynamically changing mechanical boundary conditions, rather than a fixed property of the tissue itself (Emig et al., 2021). Reproducing this interaction in *ex vivo* cardiac tissues requires more than measuring contractile force or muscle shortening under a single condition but also requires the ability to define and quantify the mechanical conditions under which that force is generated. Measurements combined with dynamic control of the mechanical environment of the tissue would provide a substantially more complete description of tissue function than any single measurement obtained under one fixed condition. Such an approach makes it possible to measure force and displacement directly and to derive shortening velocity and mechanical power from them across a range of external loading conditions applicable for *ex vivo* tissue maturation, disease modelling and pharmacological testing.

A range of experimental systems has enabled mechanical loading and functional measurements in engineered cardiac tissues, using approaches such as compliant tissue supports, mechanical actuators, magnetic fields, flexible probes, and piezoelectric actuation (Bliley et al., 2021; Helfer et al., 2025; Leonard et al., 2018; Li et al., 2023; Rodriguez et al., 2019; Sidorov et al., 2017; Xu et al., 2015). These approaches have firmly established mechanical loading as an important variable in engineered cardiac tissue culture and have enabled investigation of how mechanical conditions influence tissue function and maturation (Mannhardt et al., 2019; Schroer et al., 2017). However, most of these systems impose a fixed preload or afterload during culture. Those that vary the load change only afterload (Li et al., 2023; Mannhardt et al., 2019; Rodriguez et al., 2019) or both loads in manual steps (Helfer et al., 2025), usually with image-based force estimation, and to our knowledge none permit these mechanical conditions to be controlled independently and changed at any moment during a single, ongoing experiment without interrupting it. This central limitation is pivotal since the mechanical environment of the beating heart is intrinsically dynamic, with preload and afterload changing throughout the cardiac cycle, and their interaction with active contraction ultimately determines myocardial force generation, shortening, and power output. Moreover, contraction is often reported only as displacement of the tissue or of the post it pulls on, although how far a muscle shortens depends on the load it works against, which is rarely controlled or measured. Interpreting contraction in physical terms requires known loads and measurement of absolute force, shortening velocity and power. A system capable of altering these mechanical boundary conditions during culture would therefore offer an experimental capability fundamentally different from simply culturing tissue under a single defined mechanical condition (Cano-Jorge et al., 2025).

Here, to address this methodological gap we introduce an electromagnetic chamber (EMC) for muscle tissue culturing and biophysical characterization with electromagnetic actuators coupled with magnetic field sensor-based tissue movement detection. The system enables programmable time-resolved characterization of active and passive mechanical properties of the tissues under standard culture conditions. Electromagnetic actuation provides independent and rapid control of preload and afterload, allowing the mechanical boundary conditions of the tissue to be changed reversibly on demand, without physical manipulation of the tissue or interruption of culture. Simultaneous quantification of applied force and tissue displacement in turn provides a direct measurement of absolute contractile force from which shortening velocity, contraction and relaxation velocities, and mechanical power can all be derived. Controlled switching between preload and afterload control during culture enables rapid transitions between auxotonic and isometric contraction, allowing the predominant energy output of the muscle to be shifted, on demand, between mechanical work and heat production. The same integrated system also permits controlled mechanical perturbation for longitudinal characterization of passive tissue properties, including stiffness and viscoelastic behaviour. This is complemented by an integrated analytical framework that extracts quantitative descriptors of tissue mechanics and contractile performance from continuously acquired force and displacement signals, enabling analysis of the dynamic relationship between mechanical loading and tissue function. This framework extends engineered cardiac tissue models beyond conventional endpoint measurements and offers a platform for investigating the coupled active and passive mechanics of cardiac muscle suitable for a wide range of applications.

## Methods

### Induced pluripotent stem cell culture and differentiation into cardiomyocytes

Human induced pluripotent stem cells (hiPSCs) (ChIPSC22, Takara Bio) were maintained on dishes coated with Matrigel (Corning) in maintenance medium (E8 medium (Gibco) supplemented with 0.5 % pencillin-streptomycin (P/S, Gibco)). For differentiation to cardiomyocytes, hiPSCs were dissociated with EDTA and replated to Matrigel-coated culture plates in maintenance medium supplemented with Rho Kinase inhibitor Y-27632 (Selleckchem). After growing to confluency, the medium was changed to differentiation medium (RPMI 1640 (Gibco) with 0.1 % bovine serum albumin (Sigma), 0.033 mM ascorbic acid (Sigma) and 1 % P/S) supplemented with 6 µM CHIR99021 (Tocris). After 24 hours, differentiation medium without chemical stimulation was added and cells were cultured in this medium for 48 hours. Subsequently, cells were cultured in differentiation medium containing 2 µM Wnt-C59 (Tocris) for 48 hours, followed by differentiation medium without supplementation until cell dissociation. Cardiomyocytes were used for production of engineered heart tissues (EHT) approximately one week after spontaneous cardiomyocyte contractile activity appeared in the differentiating cells.

### Generation and culture of EHTs

Differentiated cells were dissociated into single cells in a buffer solution (NaCl 100 mM, KH2PO4 1.2 mM, MgSO4 4 mM, HEPES 10 mM, Taurine 50 mM, Glucose 20 mM, KCl 10 mM) containing 2 mg/mL type II collagenase (Worthington) and 2 mg/mL pancreatin (Sigma-Aldrich). 1.5 million cells were suspended into 47 µL of DMEM (1 % P/S) and mixed with 50 µL of solution containing 10 mg/mL fibrinogen (Sigma), 200 µL/mL Matrigel and 10 µM Y-27632 in DMEM (1 % P/S). The mixture was combined with 3 µL of 1000 U/mL thrombin (Sigma) and quickly pipetted into an agarose mold where the poles of the custom-made racks (Fig. 1A, Fig. S1B) were positioned so that the fibrin-based tissue formed between the ends of the poles. EHTs were incubated in incubator (+37 °C) for 80 minutes and 500 µl of DMEM (1 % P/S) was added. EHTs were incubated for 10 minutes with DMEM and carefully moved to a custom-made holder on a well of 6-well-plate containing 4 ml tissue medium (DMEM with 0.1 % bovine serum albumin, 0.033 mM ascorbic acid, 10 µg/ml insulin (Thermo Scientific), 2 mg/mL aminocaproic acid (Sigma) and 1 % P/S). Maintenance and long-term recordings of EHTs were performed in the tissue medium. Long-term monitoring was initiated at the day when synchronous contractile activity was detected.

**Figure 1.**
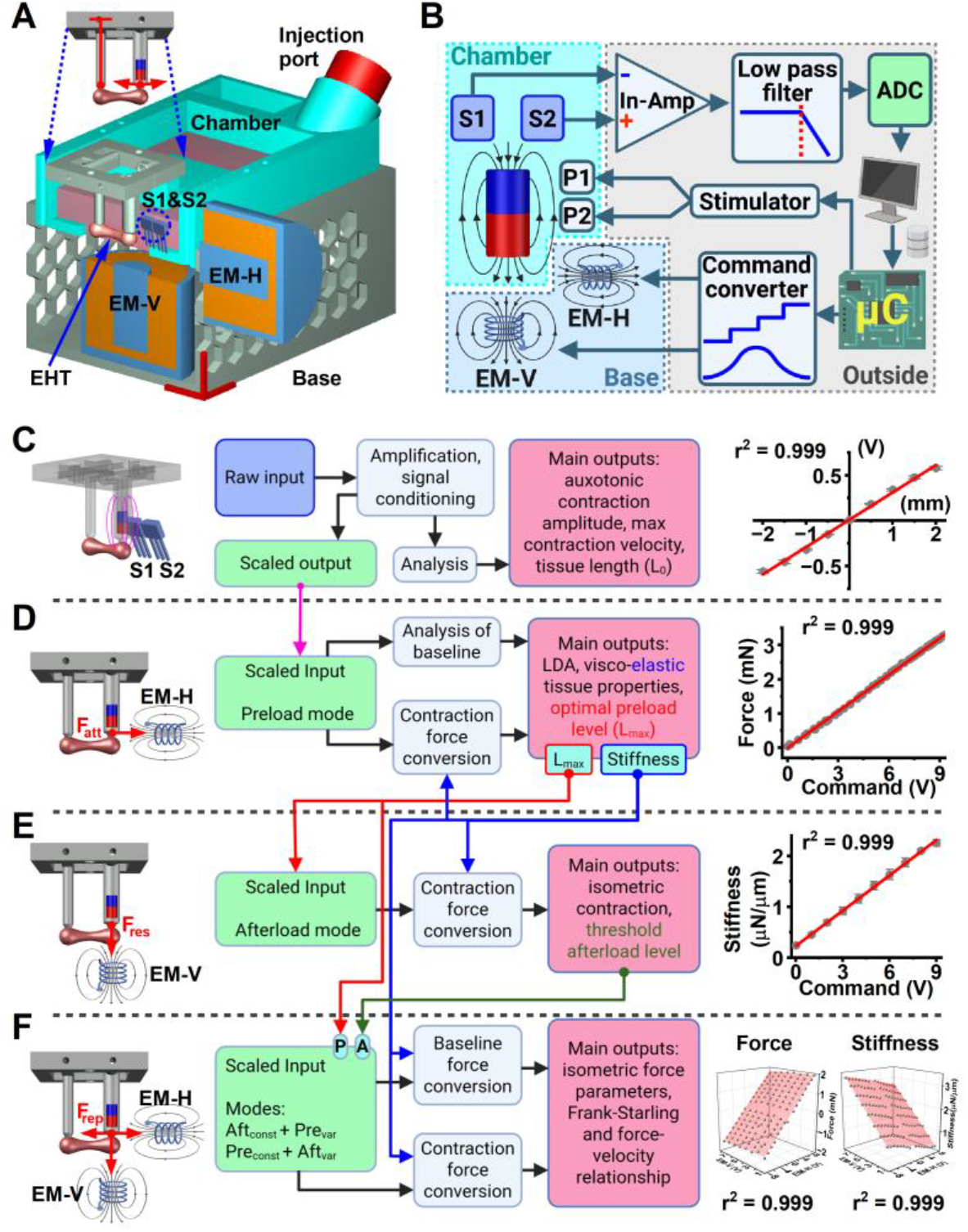
Principles of the magnetic-based control and recording system. **(A)** An overview of the magnetic-based EHT recording and mechanical loading system design in addition to 3D schematic showing detachable tissue rack, main chamber and base with electromagnetic actuators. The system consists of a rack (upper; left post is fixed and right one with magnet is movable), culture chamber with sensors (middle - light blue; EHT – heart tissue, S1CS2 – sensors; pink - culture medium) and base with electromagnetic actuators (lower grey; EM-V – vertical axis and EM-H – horizontal axis electromagnetic actuators; cross-section at the middle of the electromagnet for better visualization; red XYZ scale bars = 10mm). **(B)** Schematic representation of system components (blue/red cylinder - a magnet embedded into post, In-Amp - instrumentation differential amplifier, lowpass Butterworth filter (8^th^ order), ADC – Analog-to-Digital-converter, µC – microcomputer, PWM-to-DC command converter and electromagnets, P1CP2 – carbon pacing electrodes). **(C-D)** Summary of utilization of the magnetic field approach for measurement and manipulation of EHT (for middle part see details in text). **(C)** Schematic of magnetic field position sensing (left), and the sensor’s differential voltage to displacement conversion (right). **(D)** Application of stretching force to EHT (left), and EM-H voltage to magnetic attraction force (F_att_) conversion. **(E)** The approach for applying magnetic-induced afterload (left), and EM-V voltage to restoring force (F_res_) conversion (right; in stiffness units). **(F)** Design of coupled EM actuators for complete position and force control (left), and plane fittings (right; see large scale 3D plots in Fig. S3C) of calibration data obtained from simultaneous actions of EM-V and EM-H electromagnet actuators (fitted separately for force (stretching force) and stiffness (elastic load)). In comparison with **D**, the repelling force is also introduced (F_rep_), thus the Z-axis covers negative and positive force values). Red arrows demonstrate the directions of accessible vectors of applied forces (F_res_ – restoring, F_att_ / F_rep_ attracting/repelling forces). Red lines in **CDE -** linear fitting and red surface is the plane fitting in **F** of calibration data.

### Measurement of contractile force by force transducer

Direct force measurements were carried out under a confocal inverted microscope (FluoView 1000; Olympus, Tokyo, Japan) in a custom-made preheated perfusion chamber equipped with an isometric (IM) force transducer (HSE-HA F10 type 375), remote controlled motor lever and pacing setup. Experiments were done under constant perfusion of pH-adjusted and perfused DMEM (61965026 Gibco, pH 7.4, bubbled with 95% O_2_, 5% CO_2_) with constant electrical pacing. The force recordings were performed from slack length (L_0_) to force of maximal force development (100% length; L_max_). To measure the maximum force developed by EHTs, a custom stretch protocol was applied to record the Frank-Starling phenomenon. EHTs were stretched by a remotely controlled motor lever with a stepwise protocol at a speed of 60 µm/min, until the stretching developed no more force, and the maximum force was achieved. Analysis of the recordings was performed with ClampFit 10 (Molecular Devices, USA).

### Culture chamber and tissue rack of ElectroMagnetic Chamber (EMC)

The parts of the chamber and rack (Fig. S1) were printed with Flash Forge Creator 3 printer (FlashForge, China). Parts were printed with carbon-filled polyamide (3D filament/thermoplastic PA4035 CF, OD 1.75mm, Jabil Inc., St. Petersburg, USA), which has suitable mechanical properties and can be autoclaved. One post of the tissue rack has a stopper, and another one can be moved (Fig. 1A). A cylindrical, high temperature grade neodymium magnet (N45) covered by nickel and shielded with epoxy was inserted into tissue posts at one end and firmly sealed by PDMS. None of the electronic parts of chamber or any of the metallic components are in contact with the culture medium, and the inner wall of the chamber is covered with PDMS (Supplementary material).

### Electrical components and hardware

The EMC system consists of magnetic field sensors, amplifier, analog-to-digital converter, PC, microcontroller, command converter (Fig. S2E) and two electromagnets (EMs; Fig. 1A,B). The sensors are located in the chamber, the EMs in the chamber base, and all other components are outside of the incubator (Fig. 1B). To track EHT shortening/length, two high accuracy linear Hall Effect Sensor (A1369EUA-24-T, Allegro MicroSystems) are connected to a custom-made low-noise amplifier with active filter (Fig. S2A-C). The amplifier utilizes a precision instrumentation differential amplifier AD8222 (Analog Devices Inc., Wilmington, USA) and 8^th^-order lowpass switched-capacitor filters MAX291 (Analog Devices Inc., Wilmington, USA). The signal was low-pass filtered at 30 Hz, well above the frequency below which 90% of the signal power lies (about 5 Hz; Fig. S2D) and was further smoothed with a Savitzky–Golay filter (second order, 30 points) before velocity and power were calculated. After filtering, the displacement detection limit (SNR = 5) was about 2 µm (Fig. S2D). Data acquisition was performed with an analog to digital converter NI PCI-6052 (National Instruments, Austin, USA). For data logging and pre-processing, we used Strathclyde Electrophysiology Software (University of Strathclyde, Scotland; WinWCP5.8 and WinEDR4.0). This software was also used to generate voltage commands to EM-actuators during recording.

A single-board microcomputer (Arduino UNO) was used to control the multichannel electrical stimulator and electromagnet actuators. Digital commands from computers are passed to the microcomputer and sent to multichannel Arduino-compatible I2C Shields (Elecrow ARS27361T). This device can generate single square pulses and/or a series of them (pulse-width-modulated - PWM) at relatively high current (2A) and support bipolar applications. Outputs of these devices are directly connected to pacing carbon electrodes or through a custom-made command converter (PWM to DC converter; Fig. S2E, Supplementary material) to control electromagnets (50N). Firmware for the Arduino UNO microcontroller was written using Arduino IDE2.0, and Processing4 was used as a user interface.

### Sensor calibration and displacement to force conversion

To convert the raw electrical signal from magnetic sensors to displacement, the rack with magnetic posts was placed into a modified chamber equipped with a plastic screw (pitch 500µm) which pulls/pushes the post in the horizontal plane within a range of ±2000µm, and the electrical signal after every full screw turn was recorded and the data was linearly fitted (Fig. 1C). The obtained scaling factor was used in recording software to get scaled displacement output (Fig. 1C). To perform EM voltage to force and stiffness calibration of the EMC system, the tissue rack was placed into an altered EMC chamber and the post with the magnet was attached to a pre-calibrated force transducer via rotary joint and stiff bar (Fig. S3A). Axial movements were applied directly to the transducer within a range of ±2000µm. EM-V and EM-H voltages were applied simultaneously with a 1V step, the EM-V voltage range was up to 8V, and the EM-H range was from -8V to +8V to cover all possible voltage combinations (Fig. S3B,C). Force, displacement and EM voltages were recorded simultaneously. The displacement and force data obtained from every single EM-V/EM-H voltage combination were fitted (Fig. S3C,D) with a simple linear regression function:

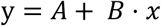

where: *A* – offset (applied force), *B* – slope (system stiffness), *y* – recorded force and *x* - applied displacement. Applied force (constant A) and stiffness (constant B) were separately plotted against corresponding EM-V/EM-H voltages, and a multiple linear regression function was used (Fig. S3C,D and Fig. 1F):

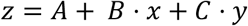

where:

- independent parameters: *x* is EM-V voltage, and *y* is EM-H voltage
- dependent parameters for applied force *A* – offset, *B* – force caused by EM-V and *C* – force caused by EM-H
- dependent parameters for stiffening *A* – offset, *B* – stiffening caused by EM-V and *C* – stiffening caused by EM-H.

The resulting formula for force conversion (Fig. S3E) with the obtained fitting constant is in the following form:

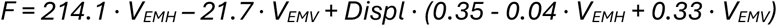

where: *Displ* is tissue displacement, *V_EMH_* is voltage at EM-H and *V_EMV_* is voltage at EM-V.

### Data analysis

To convert displacement to force Hook’s law was utilized:

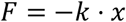

where: *k* – coefficient of stiffness and *x* is displacement. After applying Hook’s law to the EMC-system the formula is in the following form:

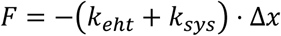

Or for instantaneous force:

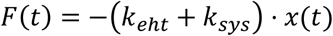

where: *k_e_*_ℎ*t*_ – coefficient of stiffness of EHT and *k_sys_* - coefficient of stiffness of the EMC-system obtained in calibration experiments, *x* is displacement (Fig. 2C), and *F* – force (Fig. 2D).

**Figure 2.**
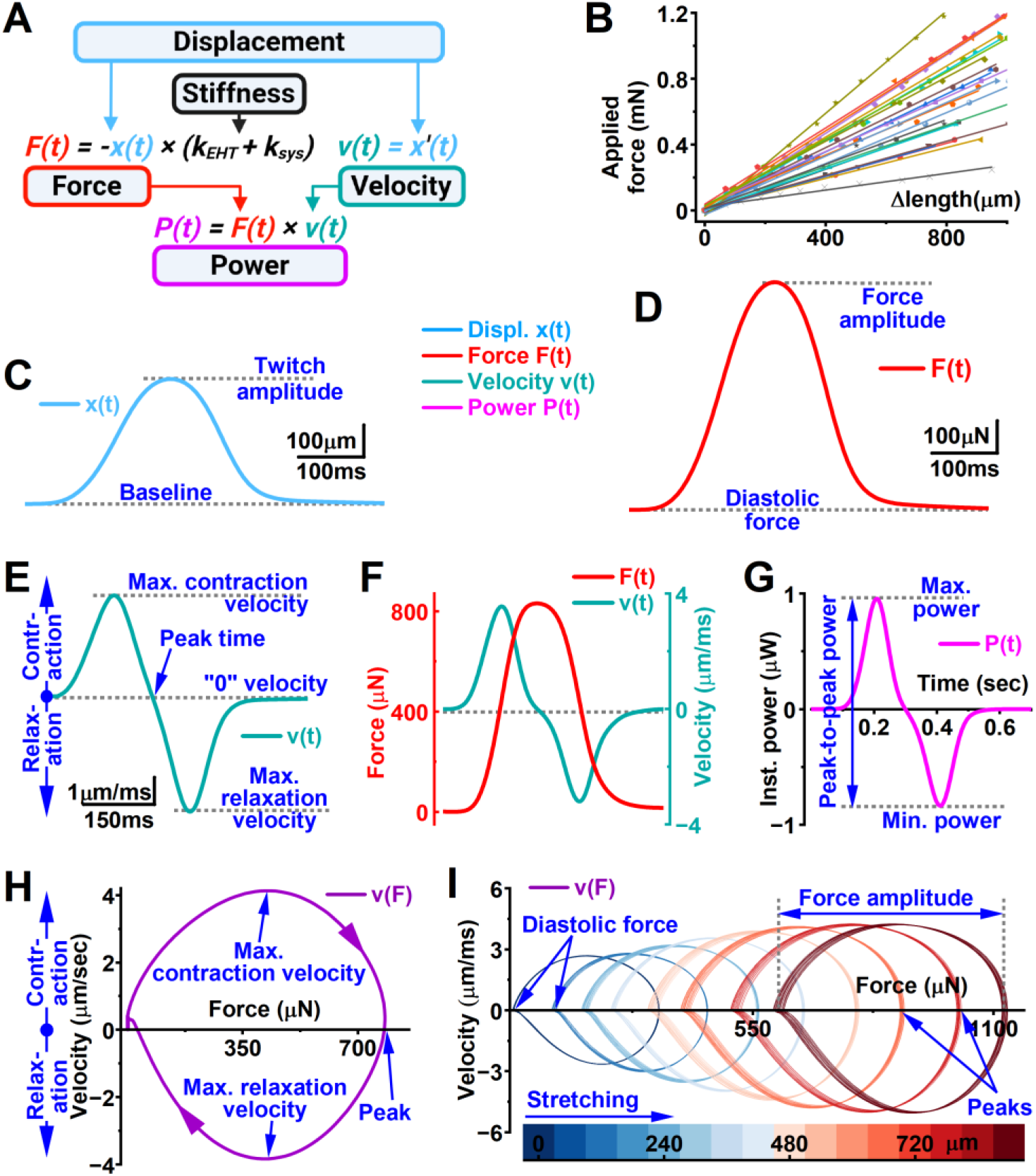
Interrogation of force, velocity and power from muscle displacement. **(A)** Flowchart and formulas used for the calculation of derived contraction parameters. **(B)** Estimation of tissue stiffness (k_EHT_; n = 23). **(C)** Representative EHT contraction trace, and **(D)** the contraction force trace obtained by converting displacement to force using tissue and system stiffness (k_sys_; note same size of scale bar in C and D). **(E)** Contraction velocity trace as a first derivative of displacement. **(FG)** An example of estimation of mechanical power. **(F)** Overlapping contraction force and velocity representative traces, and **(G)** instantaneous power calculated by multiplying contraction force and velocity. **(H)** The FV-loop (force-velocity phase plot) shows dynamic changes in force and velocity during one contraction-relaxation cycle (arrows on loop are direction of contraction), and **(I)** the FV-loop trajectory representing changes in velocities, diastolic forces and contraction forces upon EHT stretching (line colors of loops correspond to EHT Δlength color scale bar).

**Figure 3.**
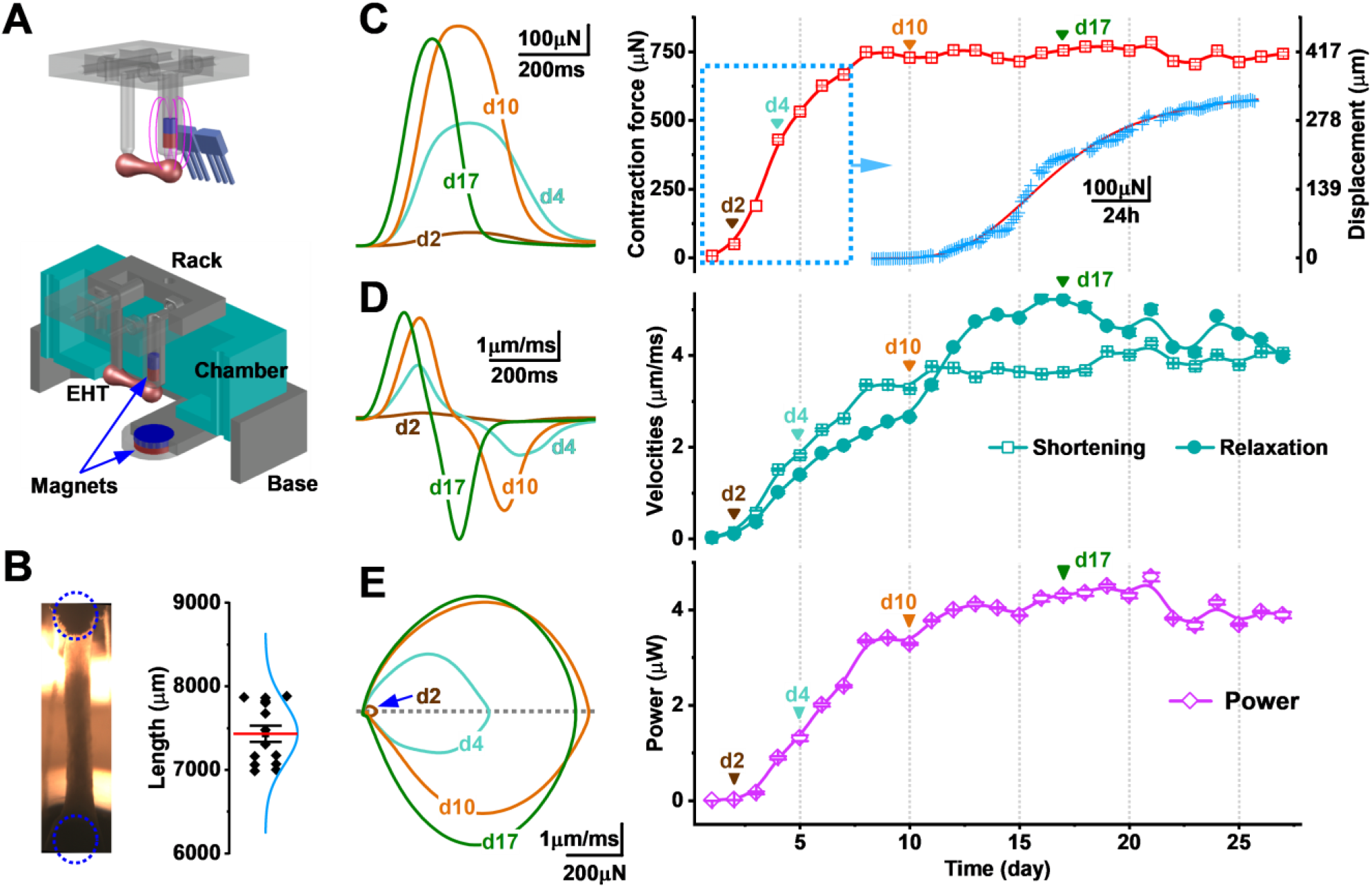
Continuous EHT contraction parameter monitoring. **(A)** 3D schematics of the use of magnet and magnetic field sensors to determine post position, and the permanent magnet base for long term EHT afterload (cross-section for better visualization). **(B)** EHT photo and tissue length measurement by sensing the magnetic post position (n = 14). **(C-E)** Representative data from long term monitoring of changes in EHT twitch parameters (note: data points are mean values collected during the day). **(C)** Representative twitch traces (left) obtained at different days (d2-d17 - time points), and twitch amplitudes (right; note: left Y-axis for force and right Y-axis for displacement; inset blue square - force amplitude at one-hour resolution). **(D)** Representative contraction traces obtained at different time points (left), and quantification of contraction velocities (right). **(E)** Representative FV-loops (left) and quantification of contraction power (right).

To get instantaneous velocity, we differentiated displacement (Fig. 2C,E):

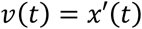

where: *v* – velocity, and *x* is displacement.

From instantaneous velocity, the maximal and minimal values were measured to obtain contraction and relaxation velocities, correspondingly (Fig. 2E). The formula for instantaneous mechanical power is:

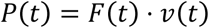

and mechanical power of contraction was assessed as peak-to-peak power (Fig. 2G):

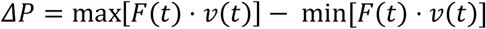

where: *P* - power, *F* – force and *v* – velocity.

### Application of mechanical load

To determine the afterload threshold, afterload was progressively increased in 0.4 µN/µm increments to identify the transition from auxotonic to isometric contraction (Fig. 5E,F). The afterload threshold was defined as the minimum afterload required at which displacement was less than 5% of the muscle length. To investigate the Frank–Starling mechanism, preload was increased stepwise in 20-µN increments, typically up to the preload corresponding to the maximum contraction amplitude (L_max_) to induce length-dependent activation (LDA) under conditions without (Fig. 4A) and with threshold afterload (Fig. 6A). In some experiments, the preload range was extended to characterize the descending limb of the force-length relationship (Fig. 4C-E).

**Figure 4.**
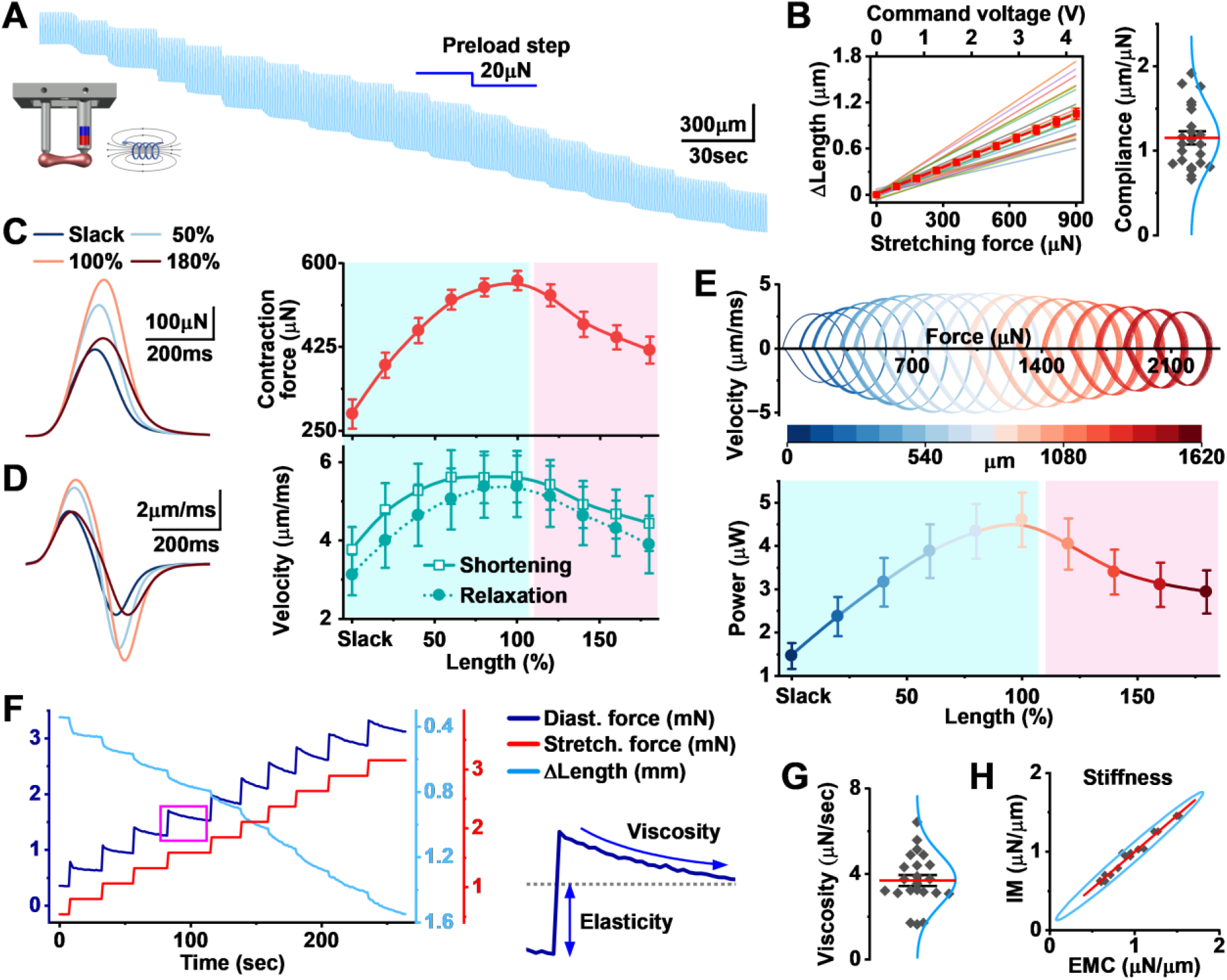
Application of preload to EHTs and estimation of passive viscoelastic tissue properties. **(A)** Representative trace demonstrating the effects of EHT stretching on twitch amplitudes and length (stretching applied in stepwise, one step is 20 µN). **(B)** Effect of stretching force on tissue length (left; transparent lines show the linear fitting from individual EHTs and bold red = mean), and estimation of force-dependent EHT elongation (right; compliance, red = mean, blue = distribution, n = 23). **(C)** Representative traces showing changes in EHT contraction force (left), and estimation of length-dependence of contraction force (light blue background – ascending limb corresponds to EHT length from slack to L_max_ and pink background color represent the descending limb). **(D)** Representative traces of contraction velocity (left) and quantification of shortening and relaxation velocities (right). **(E)** Representative FV-loop dynamic trajectory (upper) and (lower) assessment of changes in power (traces, line and symbol colors correspond to EHT Δlength color scale bar; C-E n = 14). **(F)** Representative traces of the response of tissue diastolic force and length to the application of stretching force (right: diagram showing viscosity and elasticity estimation). **(G)** Evaluation of viscosity (red = mean and blue = distribution, n = 23). **(H)** Results of pairwise comparison of EHT stiffness assessed by the system (EMC) and obtained in isometric (IM) force measurement (ratio = 0.98±0.01; red line shows linear fitting, blue ellipse is confidence level 95%, Pearson correlation = 0.99, n = 17 pairs).

**Figure 5.**
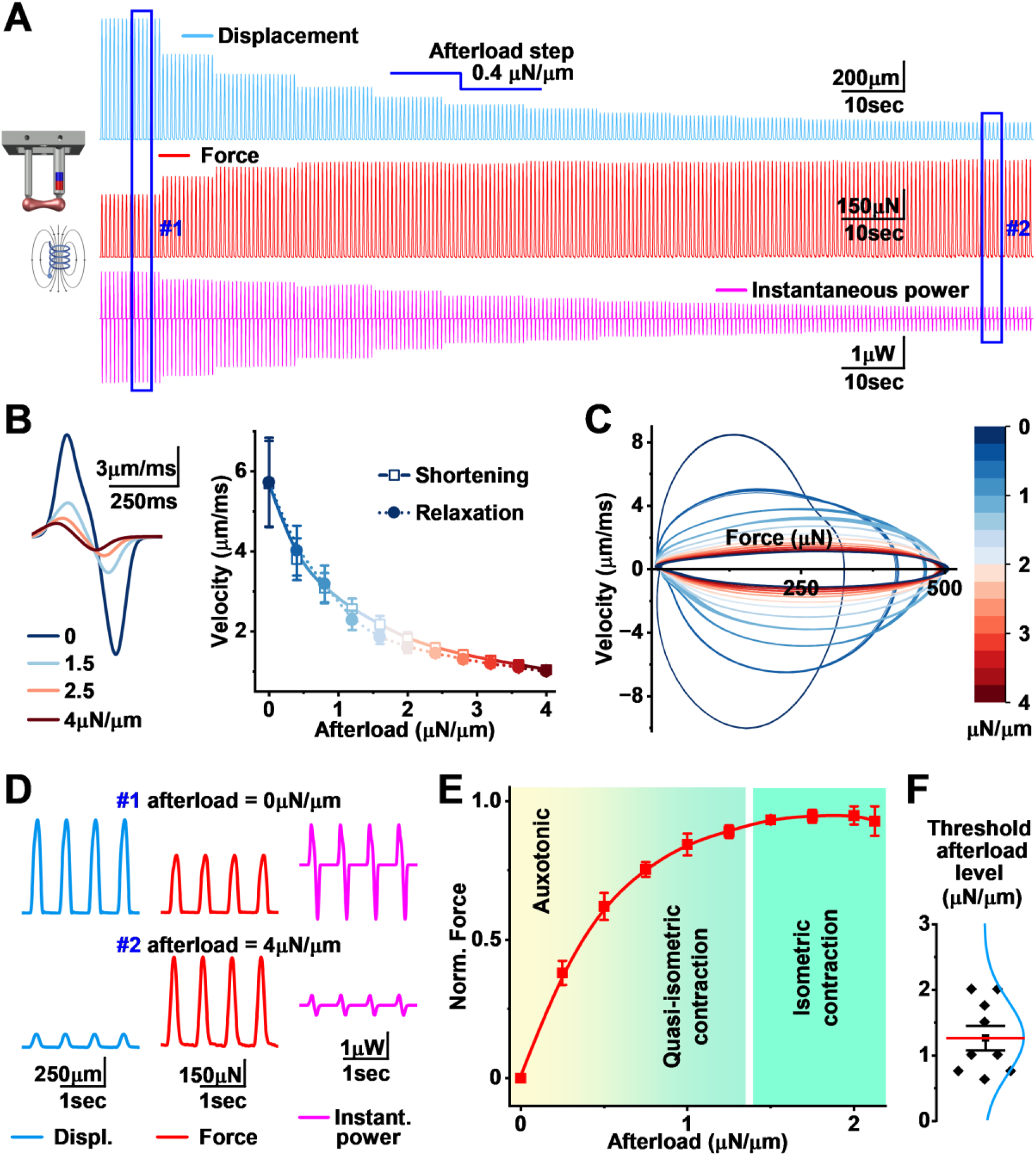
Application of afterload to EHTs and evaluation of isometric afterload level. **(A)** Representative traces showing changes in shortening amplitudes (upper), force (middle) and instantaneous power (lower) at different levels of afterload (note: afterload level shown in stiffness units)**. (B)** Representative traces of contraction velocity (left), and quantification of shortening and relaxation velocities (right) at different levels of afterload (line and symbol colors correspond to afterload level, color scale bar shown in C; n = 11). **(C)** Representative FV-loop trajectory demonstrating changes in contraction force, shortening and relaxation velocities. **(D)** The representative traces of contraction, force and velocity obtained from regions #1 and #2 shown in A, demonstrating the relationship between shortening, force, velocity and afterload levels. **(E)** The normalized forces at different afterload levels showing the relationship between afterload and contraction type. **(F)** Quantification of threshold afterload level for individual tissues (red line is mean and blue line is distribution, (F) n = 9).

**Figure 6.**
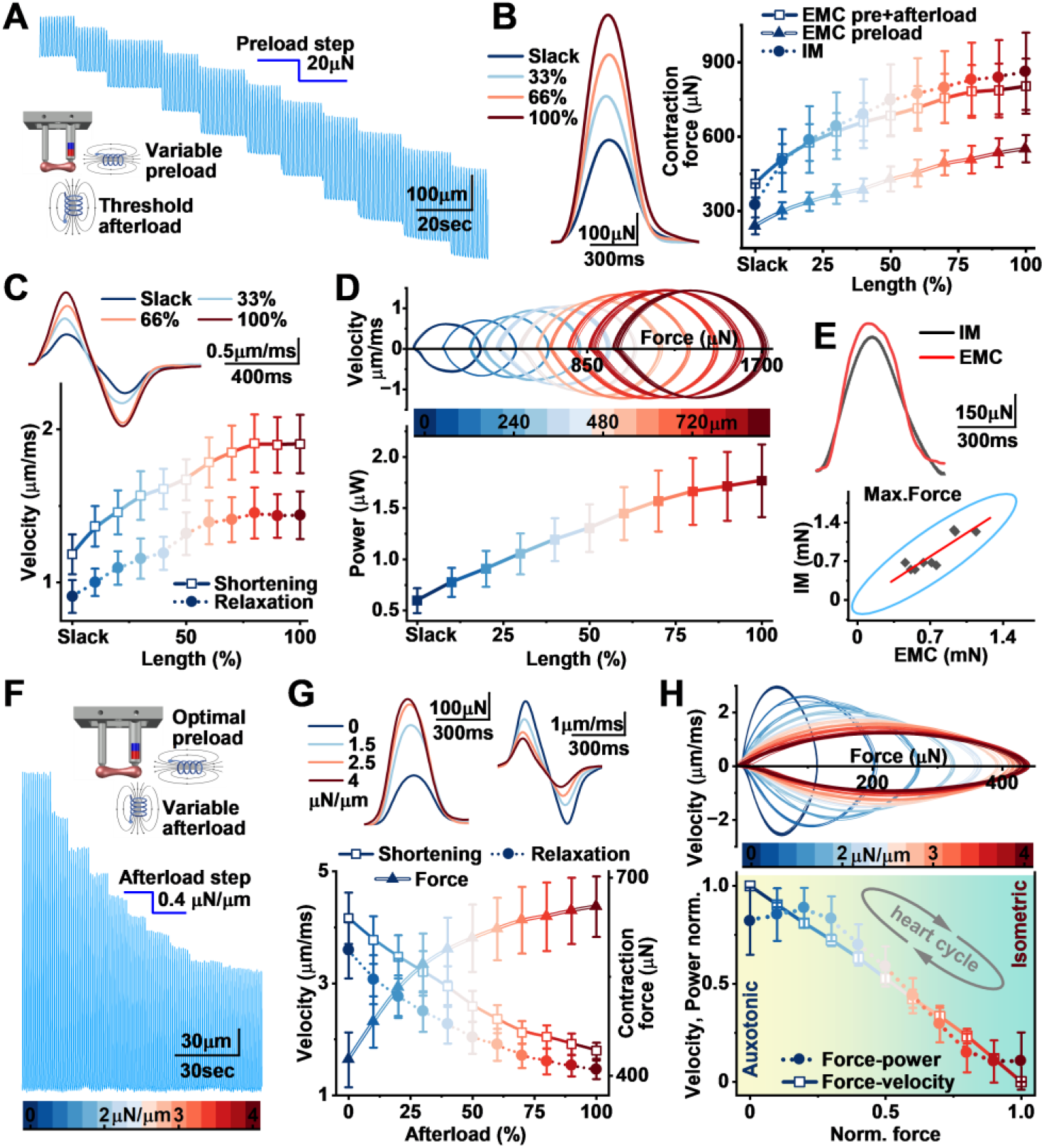
Muscle performance at varying combinations of preload and afterload (A-E) Length-force relationship. **(A)** Representative contraction trace showing changes in twitch amplitude and baseline force at different levels of preload and constant threshold afterload. **(B)** Representative contraction traces (left) demonstrating changes in force amplitude at slack, 33, 66 and 100% of L_max_, and Frank-Starling curves from preload experiments (EMC preload), preload at isometric afterload level (EMC pre+afterload) and isometric (IM) force measurement (n = 5 pairs). **(C)** Representative traces of contraction velocity (upper) and quantification of shortening and relaxation velocities (lower) at different tissue lengths (line and symbol colors correspond to EHT Δlength, color scale bar shown in D). **(D)** Representative FV-loop trajectory (upper) at different EHT lengths, and estimation of length dependent changes in contraction power (lower; C-D n = 10). **(E)** Representative force traces (left) at L_max_ recorded by the isometric force transducer (IM) and by the EMC system (EMC), and pairwise comparison of forces (right) obtained at L_max_ using the isometric force transducer and the EMC system at threshold afterload level (ratio = 0.93±0.06; red line is linear fitting, blue ellipse is confidence level 95%, Pearson correlation coefficient, r = 0.88, n = 9 pairs). **(F-H)** Force-velocity and force-power relationships. **(F)** Representative contraction trace showing changes in twitch amplitude at different levels of afterload (color scale bar) at constant optimal preload. **(G)** Representative traces of contraction force and velocity (upper), and load-dependent changes in force and velocity (x-axis: 0% – without load and 100% is load at F_max_). **(H)** Representative FV-loop dynamic trajectory (upper) at different EHT afterload (line and symbol colors correspond to EHT afterload, color scale bar), and estimation of force-velocity and force-power relationships (lower; G-H n = 9).

### Statistics

Experimental data were analyzed in OriginLab Pro 10 (OriginLab Co., Northampton, MA, USA). Results are presented as mean ± SEM. To assess the reliability of force measurement systems, measurements obtained using the EMC system and an isometric force transducer were compared using Pearson’s correlation coefficient with a 95% confidence and the mean ratio ± SEM of measurements obtained using the two methods.

## Results

### General design and analytic output of the ElectroMagnetic Chamber (EMC)

The EMC system was designed and built to enable dynamic control and measurement of physiologically relevant active and passive mechanical forces of cardiac muscle tissue during long term culture. In this context, dynamic means that preload and afterload can be adjusted during culture and measurement without handling the tissue, as the loading of the working heart changes with filling and arterial pressure. Considering the possible engineering options, we designed this system to consist of a standalone module, rather than attempting to fit the configuration into standard culturing plates. One independent EMC module accommodates one tissue, is designed to be kept in a normal cell culture incubator and consists of three separate parts: a rack for holding the tissue, the chamber to contain the culturing solution and a base with two electromagnetic actuators (Fig. 1A, Fig. S1) or alternatively magnets with fixed field strength (Fig. 3A). In the tissue rack, one post is in a fixed position and the other, with an embedded magnet, hangs from a hinge allowing the tissue placed between the two posts to move freely upon muscle movement (Fig. 1A). The main chamber consists of a port for solution injections, carbon electrodes for stimulation, and two high accuracy linear magnetic field sensors for measuring the displacement of the moving post of the tissue rack (Fig. 1A,B). The small size of the chamber, together with a single electrical cable and a solution injection port, makes it possible to house multiple chambers in a standard incubator, while the tissue environment is controlled remotely from outside the incubator (Fig. S1F). Because each module is self-contained, it can also be used outside the incubator as a bench chamber for biophysical characterization of EHTs and, potentially, other strip-like muscle preparations, with the same control of load and the same force readout. The raw signal from magnetic field sensors is acquired by using a differential instrumentation amplifier (Fig. S2A-C) and low-pass filtered to enhance the signal-to-noise ratio (Fig. S2D). A/D conversion software is used for offset subtraction and conversion to obtain measures of tissue position and movement (Fig. 1B). A microcomputer is also used to control voltage signals transmitted to stimulation electrodes and both horizontal and vertical electromagnets that apply dynamic preload and afterload to the tissue (Fig. 1B). To account for possible technical differences in sensors and EM-actuators, each EMC unit is calibrated separately, enabling full comparability of the results between EMC units (Fig. 1C-F).

In its simplest measurement mode, the magnetic field sensors of the EMC continuously record muscle movement, and after on-line conversion of the sensor voltage outputs to the calibration curves result in measures of auxotonic contraction and tissue length (Fig. 1C). When the measurements of tissue length (L_0_) and movement are combined with the attraction force (F_att_) produced by the horizontal electromagnetic actuator (EM-H; Fig. 1D), the parameters of length-dependent activation (LDA) of the muscle, viscosity, elasticity (stiffness) and optimal preload (L_max_) levels can be obtained (Fig. 1D). The measured stiffness of each tissue is a central parameter for conversion of displacement (μm) to force (μN) and L_max_ is used as a tissue-specific scaling factor to make the functional parameters of each tissue more comparable to each other.

Applying a voltage command to the vertical electromagnet (EM-V; Fig. 1A,B) generates vertical downforce establishing the restoration force (F_res_; Fig. 1E). This force increases resistance to tissue contraction, creating elastic load i.e. afterload. Applied afterload is measured as the stiffness in the horizontal plane (perpendicular to F_res_) and linearly depends on the voltage applied to EM-V. When varied, this reveals tissue specific determinants of isometric contraction (Fig. 1E). Simultaneous modulation of after- and preload makes it possible to study the central biomechanical properties of the tissue and to better simulate the physiological mechanical environment of the working heart. In practice, combining the force vectors induced by both EM-H and EM-V enable complete control of external mechanical forces, as preload and afterload can be adjusted independently and simultaneously (Fig. 1F). Applying positive (F_att_) or negative (repelling force; F_rep_) force in the horizontal direction (EM-H) controls the stretching force and thereby tissue length, and vertical force (EM-V) sets the afterload.

### Analytic tools for force, velocity and power analysis

The EMC unit measures tissue length in real time at preloads and afterloads set by directional magnetic fields from the electromagnetic actuators. Besides allowing us to measure muscle shortening, this combination allows tissue-specific stiffness to be determined, which is needed for calculating the force of contraction (Fig. 2A) because the applied load is balanced by both active and passive tension (Emig et al., 2021). Tissue-specific stiffness was assessed by measuring the response of each EHT to a series of stepwise increases in horizontal force (preload) applied with the EM-H actuator. The relationship between applied force (diastolic force; Fig. S3E) and tissue displacement was fitted by linear regression over the range of preloads applied, and the EHT-specific slope was used in the analysis as the tissue-specific stiffness coefficient (Fig. 2B). To obtain contraction force (Fig. 2D), displacement (Fig. 2C) was multiplied by a sum of EHT stiffness and EMC system stiffness (coefficient B; Fig. S3E). To account for two central features characterizing muscle function, force and velocity (Fig. 2F), the instantaneous velocity of muscle shortening was calculated as the first derivative of displacement (Fig. 2E). The mechanical power of the muscle was estimated as the peak-to-peak value of instantaneous power, thereby incorporating both shortening and relaxation velocities (Fig. 2G). To visualize dynamic changes in force and velocity, we used a phase plot of instantaneous contraction force against velocity from a single twitch (Fig. 2H), which, when extended over multiple contractile cycles, enables plotting trajectories of force-velocity loops (FV-loop), showing changes in diastolic force, systolic force and velocity during altered tissue loading in a time resolved manner (Fig. 2I).

### Long term monitoring of contractile performance of EHTs at fixed load

For long-term monitoring, the EMC system can be also equipped with permanent magnets with calibrated field strengths (Fig. 3A). To test this passive (i.e. non-dynamic) loading, we used a base with magnets with fixed field strength (0.7µN/µm) and paired magnetic sensors (Fig. 3A, Fig. S1E). This approach provides measures of post displacement but also the absolute position of the tissue free end (post with EMC embedded magnet), thus giving an accurate measure of the unloaded length of each tissue (Fig. 3B). We used this system to record contractile activity during the 27-day long maturation of iPSC-derived EHTs. Due to the high sensitivity and signal-to-noise ratio of the magnetic field sensors (Fig. S2D), the EMC detected EHT contractions from a very early stage during EHT development, even before the individual twitches became clearly visible. With a known load, the conversion of tissue shortening to force follows Hooke’s Law (Fig. 2A,D) and tissue stiffness can be derived from the degree of length change upon fixed load (Fig. 2B). As an indication of tissue maturation, the EHT shortening amplitudes increased rapidly during the first week of culture, reaching maximal values on the 8^th^ day (Fig. 3C). Interestingly, while the increase in the amplitudes of the individual twitches plateaued after day 10, the contraction velocities increased for up to 2-3 weeks (Fig. 3D). Based on these parameters, the contractile maturation of the EHTs in these conditions was complete around 20 days of culture when a plateau in the measured parameters was reached (Fig. 3C-E). The maturation of cardiac tissues involves simultaneous changes in numerous cellular systems such as excitation-contraction coupling, calcium signaling, and contractile element composition, each of which affects tissue contraction differently. The EMC system allows us to evaluate the combined effect of these processes on the contractile performance of EHTs by considering both the velocity and the amplitude of the contraction (Fig. 2F-I). To this end, we calculated the power of individual contractions (Fig. 3E), which incorporates both shortening and relaxation velocities which seemingly parallel well with the overall functional maturation of the tissue.

### Dynamic control of preload enables quantification of the length-dependent passive and active properties of the EHTs

Dynamic control of muscle preload is a prerequisite for assessing heart tissue function as preload is the driving force of length-dependent activation (LDA) of cardiac muscle and the physiological Frank-Starling response of the whole heart. Besides cellular length dependent mechanisms, one of the central determinants of the tissue LDA is stiffness (elasticity), which contributes to how much muscle lengthens at any applied preload. The EMC system enables assessing muscle LDA, but more importantly, it also makes it possible to determine the passive properties of the tissues by determining their individual preload-length change-relationship. To assess these features, we applied stepwise preload to EHTs by using the horizontal electromagnet (EM-H) of the EMC (Fig. 4A). The results show that the relationship between stretching force and the change in the EHT length is linear in the physiological range (Fig. 4B). However, demonstrating the complex biology of the EHT model, despite identical genetics and culture conditions, the compliance varies greatly between tissues (Fig. 4B). Consideration of this variability of passive mechanical properties exemplifies how essential it is to account for measures of individual passive properties in the analysis of EHT function (Fig. 1D-F, Fig. 4F-H). Characteristic for LDA, augmented EHT preload induced parallel increases in the force, velocity and power of the contraction up till maximal contraction at optimal muscle length (Fig. 4C-E). Interestingly, upon further increase of the preload the EHTs show a decline in force development, establishing the declining limb of the force-length relationship characteristic to cardiac muscle (Layland et al., 1995).

The passive mechanical properties of cardiac tissue consist of many types of dynamic and static mechanical properties that contribute to the systolic and diastolic function of the heart (Emig et al., 2021). These properties resist changes in length and thus augment the diastolic force when the tissue is subjected to stretching force i.e. preload (Fig. 4F-H). When we utilized the EMC system to study this phenomenon more closely, we found that the baseline tension response of the tissue to a length change step had two components (Fig. 4F). Each stepwise increase in stretching force led to a lasting increase in the diastolic force, indicating that the tissue resists deformation in response to an applied force, representing the stiffness of the tissue. This response also consists of a transient increase in the diastolic force, indicating that EHTs exhibit resistance to the rate of length change, demonstrating viscosity (Fig. 4F,G). To validate our results from the EMC system, we repeated the measurements by using a conventional force transducer, and the results from these systems were very similar (Fig. 4H; IM/EMC ratio = 0.98±0.01). These experiments demonstrate that the EMC system enables accurate and physiologically relevant characterization of the length dependent properties of EHTs during culture.

### Switching the mode of contraction by dynamic control of afterload

During the work cycle of the heart, the tissue shifts from isometric force generation to generating mechanical work during auxotonic contraction. In the working heart, these modes are essentially determined by shifts between the external forces, afterload and preload. To be able to induce transitions between auxotonic and isometric contraction, we used the vertical electromagnet (EM-V) of the EMC system. By applying an increasing amount of afterload (one step is 0.4 µN/µm), the transition of the contraction mode from displacement to force generation was demonstrated (Fig. 5A). At minimal amounts of afterload, muscle shortening produces mechanical work with minimal force, but upon application of increased afterload, the force generation increases at the expense of muscle work, caused by decreased contraction velocity and consequentially decreased mechanical power (Fig. 5B,C) as well as shortening (Fig. 5D). The amount of external force required to shift from auxotonic to isometric contraction varies among EHTs (1.26±0.18 µN/µm) and is affected by tissue stiffness, but with a gradual increase in afterload it is possible to find a tissue-specific isometric threshold (Fig. 5E,F). This isometric threshold, defined here as the afterload at which shortening during the twitch reduces to less than 5% of the muscle length, was determined from the relationship between applied afterload and shortening recorded with the EMC. Because this parameter relies on the calibrated actuator load instead of the displacement-based force conversion, it provides an independent check of the latter, and it allows afterload to be expressed relative to the force capacity of each tissue when force-velocity and power relationships are compared. With the option of making the contraction isometric, it is also possible to determine the tissue- and preload specific maximal isometric force, representing the force produced by the muscle when subjected to load constraining muscle shortening to zero (P_0_) (Hill, 1938; Seow, 2013).

### Simulation of the physiological mechanical environment of cardiac contraction

During the cardiac cycle, muscle is subjected to sequential changes in tissue length and elastic load, and these establish the mechanobiological framework for the pressure-volume loop of heart contraction. In various pathological conditions, these forces are disproportionately changed, such as preload increases in heart failure (HF) upon increased volume load and afterload increases due to hypertension. To demonstrate the combined effect of these two forces on EHTs, we varied the preload to trigger LDA at fixed afterload inducing maximal isometric contraction (Fig. 6A). Upon increased preload, EHTs showed an increase in the force amplitudes (Fig. 6B), contraction velocities (Fig. 6C) and power (Fig. 6D), all of which were increased across the typical range of the LDA response. Expectedly, afterload seems to have a significant impact on LDA, as demonstrated by comparison to non-loaded (without afterload) EHTs (Fig. 6B). To validate the force recordings of the system, we repeated the measurements by using a standard isometric force transducer instead of a magnetic field detector for force recordings, and both methods produced comparable LDA (Fig. 6B) and maximal force (Fig. 6E; IM/EMC ratio = 0.93±0.06). To study the effects of variable afterload at constant preload, we first determined the optimal muscle length to produce maximal LDA by varying preload while maintaining afterload at isometric threshold i.e. afterload required to make the contraction isometric. With fixed optimal preload, a stepwise increase in the afterload (Fig. 6F) results in an increase in force which is accompanied by a decrease in contraction velocity (Fig. 6G,H). These results demonstrate that with the EMC, preload and afterload could be varied independently in the same EHTs. Together, these results show that the EMC reproduces the load-dependent behavior of cardiac muscle (force-length and force-velocity relationships) with independent, adjustable control of both loads in tissues kept in culture. This makes it possible to test, in the same tissue, how the combined preload and afterload changes common to physiological or pathological settings affect muscle function, and this lays the experimental foundation for studying long term effects of specific combinations of these stimuli.

## Discussion

In addition to traditional *ex vivo* experimental cardiac models such as cultured human heart tissues (Fischer et al., 2019; Perbellini et al., 2018; van der Geest et al., 2025) and rodent tissue preparations (Cao-Ehlker et al., 2023), human iPSC-derived engineered heart tissues (EHTs) have emerged and proven useful as preclinical platforms for drug screening (Anand et al., 2025), experimental tools for mechanistic studies (Campostrini et al., 2021) and elucidation of monogenic cardiac disease mechanisms (Carvajal-Vergara et al., 2010; Itzhaki et al., 2011; Yazawa et al., 2011). Their functional maturity is still limited (Andrysiak et al., 2021; Eder et al., 2016), and most culture systems neither reproduce the *in vivo* cardiac tissue environment nor allow physiologically relevant measurements (Cano-Jorge et al., 2025). Addressing both requires methods that simulate the *in vivo* mechanobiological environment of the developing and adult heart. Pneumatic actuators, stretchable membranes and magnetic beads can apply preload to cardiomyocytes (Cano-Jorge et al., 2025), but these do not allow independent control of both loads or direct force readout in EHTs in culture. The chamber described here provides programmable electromagnetic control of both loads, preload and afterload, which can be adjusted freely during culture. Length-dependent activation, passive stiffness and viscosity can be measured in the same chamber, which moves in-depth biophysical muscle experimentation into the culture incubator while keeping the mechanical environment under experimental control. The same chamber can also be used as a stand-alone perfusable setup for biophysical characterization of EHTs and, potentially, other strip-like muscle preparations, with the same control of load and measurement of force.

Recapitulating heart function *ex vivo* is fundamentally difficult, yet it is essential for meaningful disease modeling and translational research. The heartbeat consists of distinct phases (isovolumic relaxation, filling, isovolumic contraction and ejection), in which myocardial strain changes at different rates (Chung, 2019; Russel et al., 2009). The helical, three-dimensional architecture of the ventricular wall produces strain that varies in space and time (Sengupta et al., 2006). Bioengineered tissues and isolated muscle strips are necessarily uniaxial and cannot reproduce this complexity. However, in both isolated cardiac preparations and in the whole heart, muscle performance is determined by preload (end-diastolic fiber length), afterload (systolic wall tension) and the inotropic state of the myocardium (Braunwald et al., 1967). Passive properties, the intrinsic stress-strain relationship of the muscle arising from sarcomeric crossbridge attachment and from the extracellular matrix and non-cardiomyocytes (Crocini and Gotthardt, 2021; Emig et al., 2021) are equally central determinants of tissue behavior, as they shape how the muscle responds to load independently of active contractile state. Consequently, active and passive properties are only separable if force and movement can be measured under independently adjustable pre- and afterload conditions, and the distinction between altered contractility and altered passive mechanics can be made.

Physiologically relevant cardiac models need to account for the fact that preload determines the diastolic length, whereas afterload opposes muscle shortening. Both have unique effects on cardiac tissue function, signaling and adaptation (Pitoulis et al., 2022; Toischer et al., 2010) as well as on the maturation of engineered heart tissues (Fink et al., 2000; Leonard et al., 2018). To our knowledge, no earlier system combines independent, adjustable control of both preload and afterload with continuous, calibrated readout of force and shortening in the culture incubator (Cano-Jorge et al., 2025). In our system, both loads are set by the voltage applied to the electromagnets and can be changed at any moment during an experiment. This allows length-dependent force changes to be examined during culture rather than in separate setups outside the culture chamber (Tuomainen et al., 2026) and enables rapid switching between auxotonic and isometric contractions whenever needed. Because the loads are known, each tissue’s own passive stiffness can be measured and used for the displacement-to-force conversion, and the isometric threshold, which is defined as the afterload at which the contraction becomes isometric, can be read directly from the tissue’s response. While more sophisticated setups allow in-depth analysis of EHT mechanical properties and load-dependent contractile features (Sewanan et al., 2021), they require interrupting the culture for laborious experiments outside the culturing environment.

Magnetic position sensing has been used for continuous force recording in culture (Bielawski et al., 2016; Fischer et al., 2019; Lu et al., 2021), as in the present study, but video-based motion tracking has become the dominant approach for characterizing contractile performance of engineered cardiac tissues (Rivera-Arbelaez et al., 2022; Sala et al., 2018; Scalzo et al., 2021). In imaging-based approaches, muscle force is described as movement of the tissue, or deflection of the pillar with desired elastic properties attached to muscle which is converted to force via a beam-bending model. A recent comparison of three widely used tracking algorithms found that, applied to identical samples, they produced meaningfully divergent kinematic estimates and, in some conditions, disagreed on the direction of the measured effect (Burattini et al., 2024). This demonstrates that video analysis does not converge on a single accurate measure but instead yields algorithm-dependent estimates. Video-based data collection is further constrained by image acquisition rate, which limits temporal resolution and trades off against light exposure and data volume at higher acquisition speeds. In the EMC system, tissue displacement is measured directly and continuously via magnetic field sensing that, unlike high-pass-filtered systems (Bielawski et al., 2016), preserves the absolute post position and thus tissue length, and force is then derived from displacement using a chamber- and tissue specific stiffness relationship. Unlike the fixed material constants assumed in post-bending systems, this stiffness is measured directly across the physiological range of preloads for each individual tissue. Demonstrating the necessity of this approach, we showed that compliance varied more than two-fold between different EHTs. As the passive properties can be re-measured at will, as often as needed over the course of culture, it allows the calibration to be updated when the mechanical properties of the tissue change over time. Because measurement and actuation operate on this same fast timescale, the system could enable real-time feedback between contractile output and applied load, adjusting the mechanical boundary condition dynamically in response to the measured behavior of the tissue, a loop that does not depend on image acquisition and processing.

Though now commonplace in disease modelling and safety toxicology, there is a broad consensus that the fundamental drawback of hiPSC-derived cardiomyocytes is an immature phenotype when compared to adult counterparts (Anand et al., 2025; Eder et al., 2016). This is manifested as insufficiently developed contractile elements and Ca^2+^-release structures, in addition to drastically different electrophysiological properties (Ewoldt et al., 2025). From early heart development, mechanical loading accelerates tissue maturation (Tobita et al., 2005), which seems to be a general feature of heart development also recapitulated by the iPSC-models (Leonard et al., 2018; Yang et al., 2014). The EMC can be used to address these deficiencies in several ways. Firstly, continuous measurement of EHT force production, together with derivation of more specific maturation measures of contraction allows for determination of the time point at which maturation has been reached for individual EHTs prior to experimental perturbation. Secondly, the EMC allows for efficient development of specific protocols which utilize preload and afterload to promote tissue maturity. With the system described in the present study, the amount and timing of preload and afterload that promote maturation can be tested systematically in EHTs, potentially allowing for the generation of human cardiomyocytes which are more suitable for pharmacological studies and disease modelling. Finally, the EMC offers a streamlined system for examining the effects of these factors and optimizing the dose needed to achieve improvement in specific maturation parameters (maximum force, power, etc), with or without accompanying mechanical loading. This should allow for better identification of optimal treatment windows and interactions between biophysical stimulation and the activation or repression of molecular pathways driving the maturation process.

The EMC setup is also well-designed for pharmacological studies, including continuous measurement capabilities, possibility for pre-programming standardized protocols, and reasonable throughput, with several chambers running side by side in a standard incubator. Our methodology also offers several measurement readouts which could be utilized to assess efficacy, such as contractile force, shortening/relaxation velocity, and power as a corollary of energy consumption due to contraction.

There are some limitations associated with this work. First, this approach does not allow for loading of the tissue in a multidirectional manner, which corresponds to the mechanical stresses experienced by the ventricular wall in physiological settings. Force is calculated based on displacement, and the calculation is done assuming that there is linear elasticity with no consideration of viscoelastic properties and no change of stiffness upon tissue activation. The force measurement was validated only for the passive stiffness and isometric force at L_max_ compared to the isometric force transducer. As one EHT can be placed into each module only, the throughput is limited compared to the plate-based approach. Dynamic loading was performed during short experiments while long-term cultivation and loading of tissues were performed only with permanent magnetic fields. In addition, closed-loop control of loading based on the tissue signal has not been tested.

In conclusion, the EMC system allows preload and afterload to be controlled independently and changed at any moment during an ongoing experiment, together with continuous readout of force, velocity and power inside the culture incubator. Tissue-specific calibration of passive mechanics renders a displacement measurement into accurate measures of contraction force while considering variability between tissues, making the results comparable across tissues and experimental conditions. We expect the platform to be most useful for defining how specific loading patterns drive EHT maturation, for separating changes in contractility from changes in passive and active mechanics in disease models, and for testing drugs under physiologically relevant load.

## Authorship contribution statement

**Nikolay Naumenko:** data curation, methodology, software, hardware design, formal analysis, investigation, writing – review C editing.

**Tomi Tuomainen:** data curation, formal analysis, investigation, writing – review C editing.

**Robert S. Leigh:** data curation, formal analysis, investigation, writing – review C editing.

**Pasi Tavi:** conceptualization, data curation, formal analysis, supervision, funding acquisition, writing – original draft.

## Data availability

The data generated during this study (including CAD files, PCB layouts, firmware and parts list) are available from the corresponding author on reasonable request. including CAD files, PCB layouts, firmware and parts list

## Acknowledgements

John Dempster (Strathclyde Institute for Pharmacy C Biomedical Sciences) for the great data record and analysis software.

## Supplementary Material

**Supplementary Figure 1.**
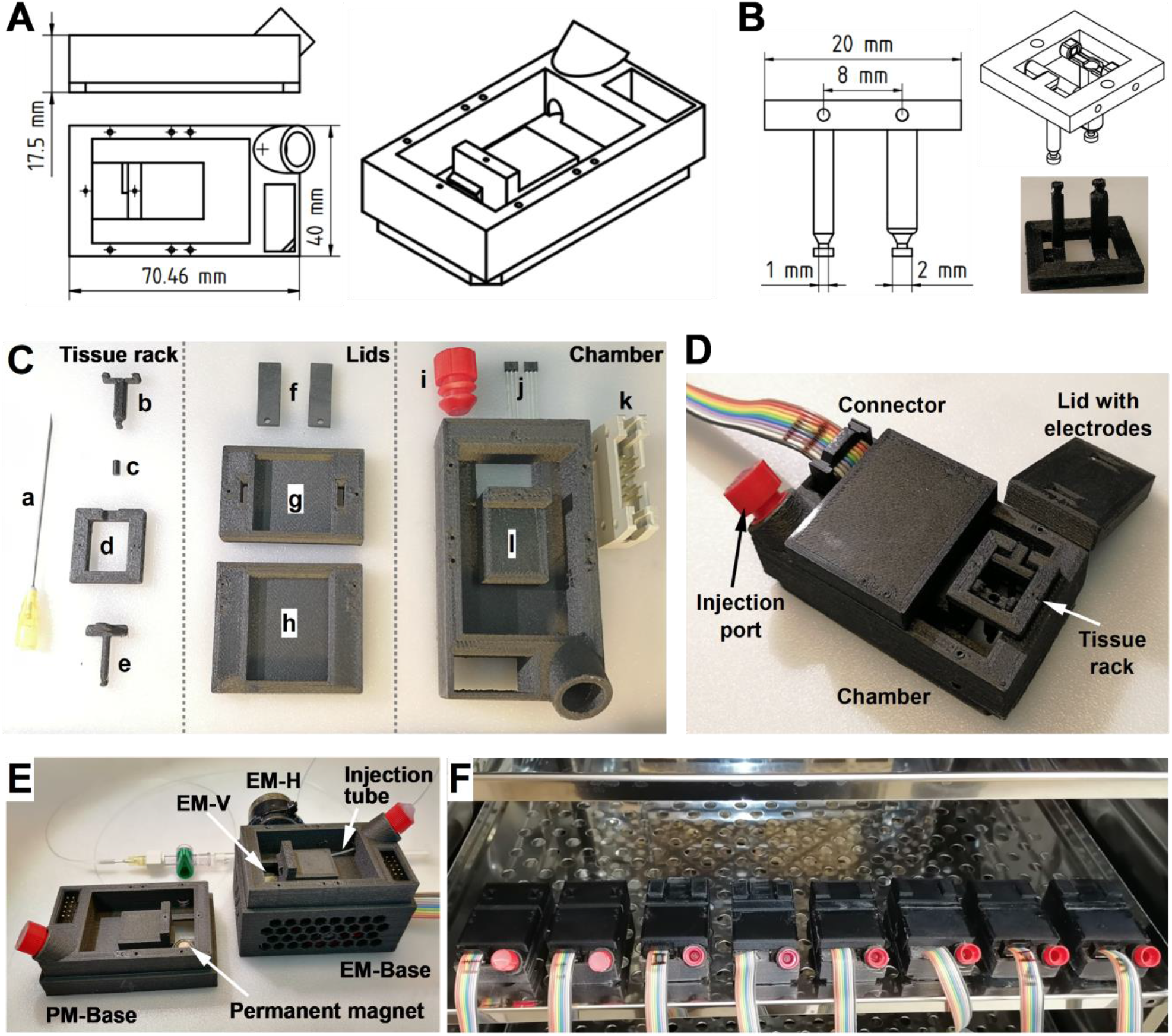
Components of EMC chamber. **(A)** Technical drawings of the EMC culture chamber and tissue rack **(B)** with dimensions. **(C)** Parts of tissue rack and culture chamber: (a) medical grade stainless needle used as a rotary axis, (b) hinged post, (c) a magnet embedded into the post, (d) tissue rack frame, (e) fixed post, (f) carbon electrodes, (g) chamber lid holding electrodes, (h) chamber lid, (i) cap of injection port, (j) magnetic field sensors, (k) connector and (l) main culture chamber. **(D)** Assembled EMC chamber. **(E)** EMC chambers on permanent magnet base (PM-Base, left), and electromagnet base (EM-Base, right). **(F)** EMC chambers in culture incubator.

**Supplementary Figure 2.**
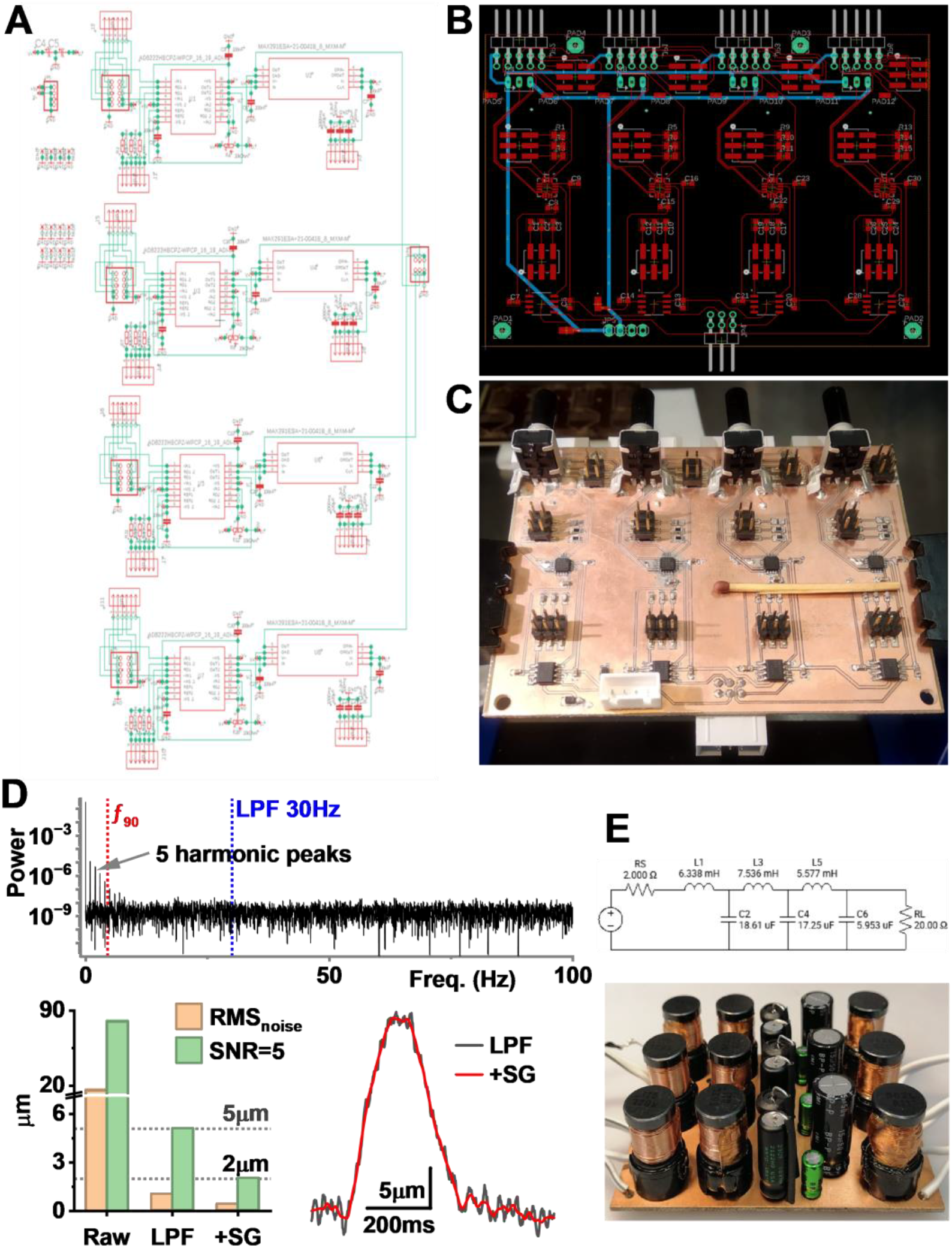
Multichannel differential amplifier with signal filter, estimation of proper filter settings, and voltage command converter. **(A)** Electrical circuit diagram of four channel differential amplifier with signal filter. **(B)** PCB layout and **(C)** realization of multichannel amplifier. **(D)** Estimation of system sensitivity. Upper: FFT power spectrum of raw signal showing 90% cumulative power frequency (red, ƒ_90_ -containing 90% of the cumulative spectral power), and settings of frequency corner of low-pass filter (LPF 30Hz). Lower left: estimation of root mean square of noise (RMS_noise_) and system sensitivity at signal-to-noise ratio = 5 (SNR=5) at different conditions: without filtering (Raw), low-pass filtered at 30Hz (LPF) and LPF combined with off-line filter (+SG; second order Savitzky-Golay). Lower right: An example of 25 μm twitch after LPF (black) and LPF combined with off-line filter (+SG). (E) Electrical circuit diagram of command converter (upper; one channel of PWM-to-DC), and realization of three channel converter (lower).

**Supplementary Figure 3.**
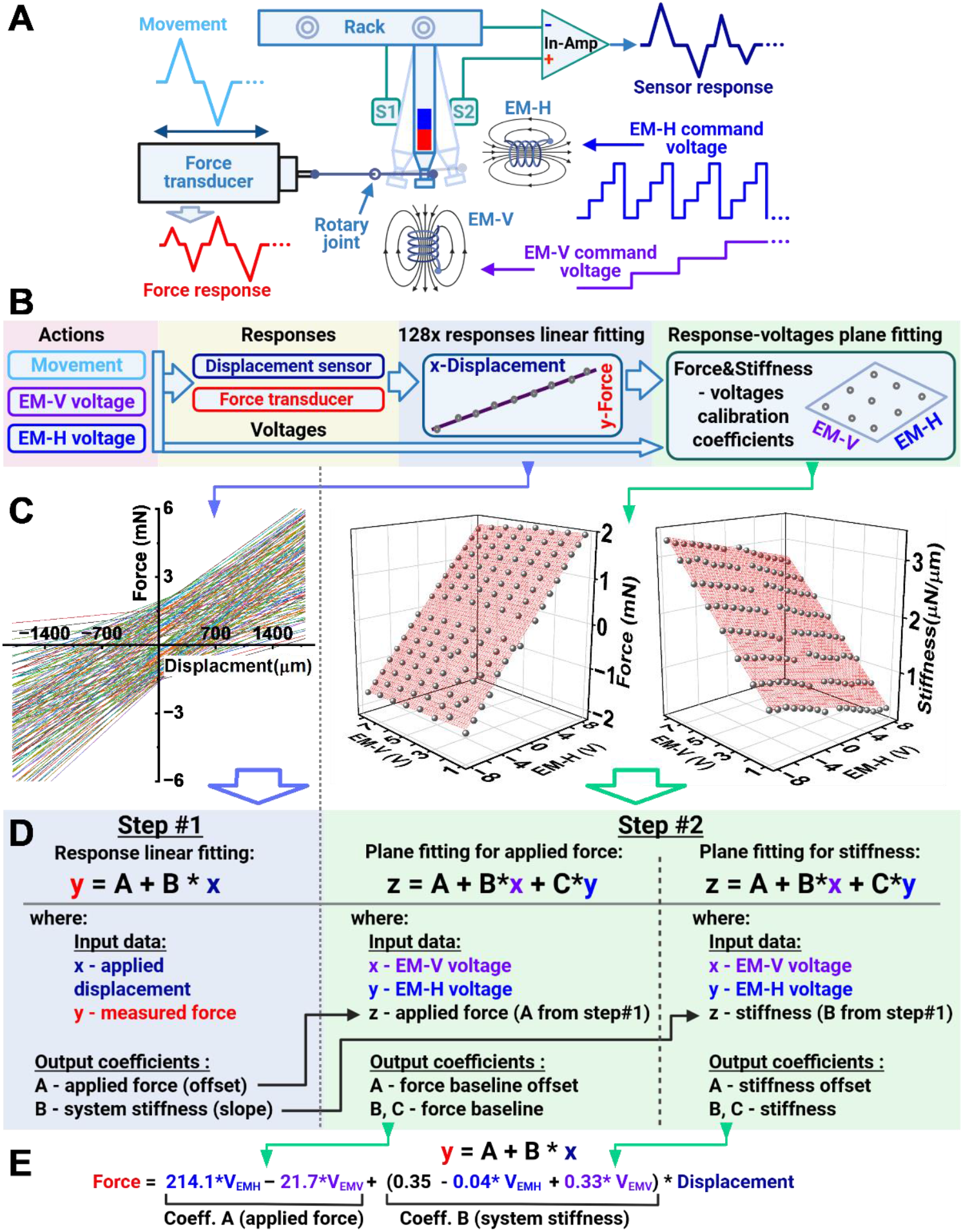
Algorithm of displacement-force-voltages to force-stiffness calibration of EMC system. **(A)** Schematic of calibration system. Tissue rack in modified chamber with pre-calibrated magnetic field sensors (S1 and S2). The rack post with the magnet is attached to a pre-calibrated force transducer via rotary joint and stiff bar. Axial movements were applied directly to the transducer within a range of ±2000µm. EM-V and EM-H voltages were applied simultaneously with 1V step, EM-V voltage up to 8V and EM-H range from -8V to +8V to cover all possible combinations (128). Force, displacement and voltages were recorded simultaneously and analyzed off-line. **(B)** The flowchart explaining the full calibration procedure from data acquisition to obtained conversion coefficients. **(C)** Displacement-force line fitting data (left; 0.9 < r^2^ < 0.999) was obtained from every single EM-V/EM-H voltage combination. Plane fittings of applied force (middle) and system stiffness (right; note: both shown on Fig. 1F for demonstration). **(D)** Explanation of the fitting steps to obtain calibration coefficients. **(E)** The final formula with calibration coefficients, coefficient A (Coeff. A) represents changes in diastolic force (applied force) and coefficient B (Coeff. B), which is system stiffness.

**Supplementary Figure 4.**
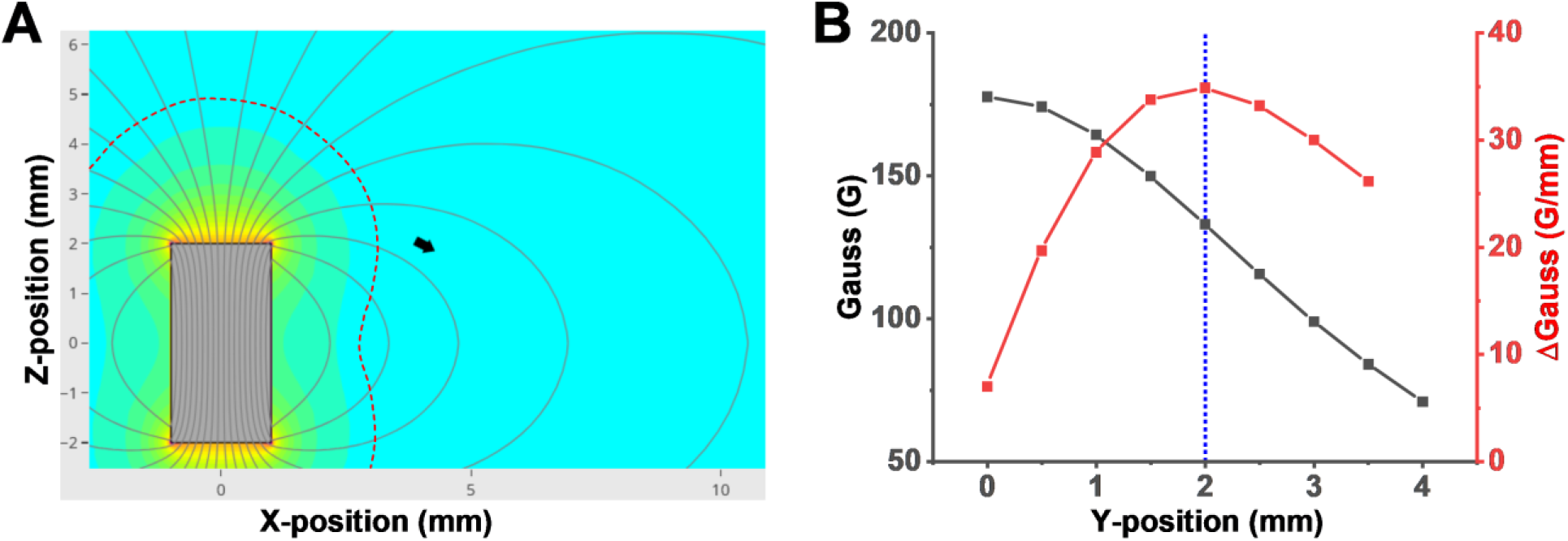
Magnet-to-Sensor optimal position calculation. **(A)** Calculation of Z-position and sensor plane angle. The black arrow is the optimal location of the sensor in the XZ plane, and the vector arrow shows the direction of magnetic field lines. The dashed line is the approximate sensor oversaturation area. **(B)** Calculation of magnetic flux density in the Y- direction (black; Y-direction is tissue rack magnetic post movements) and changes in density upon moving in the field (red), showing changes in flux density upon movements (blue dotted line is distance between centers of sensors in the EMC-system).

## Supplementary methods

### Culture chamber and tissue rack

The culture chamber is built in a modular format. The EMC chamber consists of four main parts: main chamber (Fig. S1A), tissue rack (Fig. S1B), the chamber lids (Fig. S1C) and an exchangeable/removable base (Fig. S1E). The parts were printed with Flash Forge Creator 3 printer (FlashForge, China), using carbon-filled polyamide (3D filament/thermoplastic PA4035 CF, OD 1.75mm, Jabil Inc., St. Petersburg, USA), which has suitable mechanical properties, biocompatible polyamide, and can be autoclaved. One post of the tissue rack has a stopper, and another one can be moved (Fig. 1A). A cylindrical, high temperature grade neodymium magnet (up to 150°C; material NdFeB, N45SH; OD:2mm, height: 4mm; custom ordered from https://www.supermagnete.fi/) with axial magnetization covered by nickel and shielded with PTFE epoxy (Ni-Cu-Ni-Epoxy and PDMS) is inserted into tissue posts at one end and firmly sealed by PDMS. The sensors are located on the bottom side of the chamber. The optimal position and angle of the sensors (Fig. S4AB) were calculated utilizing on-line tools (https://www.kjmagnetics.com/magnetic-field-calculator.asp - for plane and https://magnetrechner.de/en/field-as-required-in-space/disk/axially-magnetized - for in space orientation). Sensors, wires and the inner surface of the chamber were covered with PDMS so that none of the electronic parts of the chamber or any of the metallic components are in contact with the culture medium. Carbon electrodes are located in the chamber lid (Fig. S1CD; EHT technology). The assembled EMC chambers were autoclaved without changes in geometry or mechanical/other properties.

### Signal amplification and conditioning

Two high accuracy linear Hall Effect Sensors (analog output, A1369EUA-24-T, Allegro MicroSystems) were used to track EHT shortening/length. The outputs of sensors were connected to custom-made low-noise multichannel amplifier with an active filter (Fig. S2A-C). The amplifier utilizes a precision instrumentation amplifier AD8222 in differential mode (Analog Devices Inc., Wilmington, USA), gain = 4 (with optional on-board switcher to set gains: 10 and 20) and offsets the output signal. Utilizing two sensors and differential input allows us to reach high output linearity and reject common mode noise. A lowpass 8^th^-order Butterworth switched-capacitor filter MAX291 (Analog Devices Inc., Wilmington, USA) was used for signal conditioning and provides a maximally flat passband response and 48dB of noise attenuation per octave. The corner frequency was set at 30Hz (with optional on-board switcher to set f_c_ = 10 or 100Hz). To find an appropriate corner frequency, raw signals were analyzed with FFT (Fig. S2D) and to estimate noise level root mean square (RMS, a mathematical measure used to find the average magnitude of a varying quantity) was used. Ratio-to-Noise = 5 (RMS*5) was a minimal accepted level for signal detection. Since shortening velocity (first derivate of displacement) is sensitive to signal quality, to improve the accuracy an additional digital filter was applied (Fig. S2D; Savitzky–Golay, 30 points, second order). The printed circuit board of the four-channel amplifier with active filter was created in Autodesk EAGLE software.

### Command converter

A single-board microcomputer (Arduino UNO) was used to control the electromagnet actuators. The digital signal was converted to PWM signal (pulse-width modulation) by using an integrated circuit (multichannel motor driver TB6612 (I2C), Elecrow ARS27361T) followed by PWM-to-DC converter (Fig. S2E), which is a 6^th^ order Chebyshev lowpass filter (cutoff frequency is 720Hz, passband ripple is 0.1 dB), designed to pass a minimum of 2A at 12 Volts. Since TB6612 utilizes an h-bridge, it allows us to apply both attraction and repelling forces by changing voltage polarity on the electromagnet. Correspondingly, a PWM-to-DC converter was designed using bipolar capacitances. The firmware for the Arduino UNO microcontroller was written using Arduino IDE2.0, and Processing4 was used as a user interface.

